# Speech Synthesis from ECoG using Densely Connected 3D Convolutional Neural Networks

**DOI:** 10.1101/478644

**Authors:** Miguel Angrick, Christian Herff, Emily Mugler, Matthew C. Tate, Marc W. Slutzky, Dean J. Krusienski, Tanja Schultz

## Abstract

**Objective:** Direct synthesis of speech from neural signals could provide a fast and natural way of communication to people with neurological diseases. Invasively-measured brain activity (electrocorticography; ECoG) supplies the necessary temporal and spatial resolution to decode fast and complex processes such as speech production. A number of impressive advances in speech decoding using neural signals have been achieved in recent years, but the complex dynamics are still not fully understood. However, it is unlikely that simple linear models can capture the relation between neural activity and continuous spoken speech.

**Approach:** Here we show that deep neural networks can be used to map ECoG from speech production areas onto an intermediate representation of speech (logMel spectrogram). The proposed method uses a densely connected convolutional neural network topology which is well-suited to work with the small amount of data available from each participant.

**Main results:** In a study with six participants, we achieved correlations up to *r* = 0.69 between the reconstructed and original logMel spectrograms. We transfered our prediction back into an audible waveform by applying a Wavenet vocoder. The vocoder was conditioned on logMel features that harnessed a much larger, pre-existing data corpus to provide the most natural acoustic output.

**Significance:** To the best of our knowledge, this is the first time that high-quality speech has been reconstructed from neural recordings during speech production using deep neural networks.

## 1. Introduction

The ability to speak is crucial for our daily interaction. However, a number of diseases and disorders can result in a loss of this ability, for example due to severe paralysis. Various technologies have been proposed to restore the ability to speak or to provide a means of communication (see [1] for a review), including Brain Compuer Interfaces (BCIs, [2]). The most natural approach for BCIs would be to directly decode brain processes associated with speech [3]. Invasively-measured brain activity, such as ECoG is particularly well-suited for the decoding of speech processes [4, 5], as it balances a very high spatial and temporal resolution with broad coverage of the cortex.

In recent years, significant progress has been made in the decoding of speech processes from intracranial signals. The spatio-temporal dynamics of word retrieval during speech production was shown in [6]. Other approaches demonstrated that speech can be decoded from invasively measured brain activity such as words [7], phonemes [8, 9], phonetic features [10, 11], articulatory gestures [12] and continuous sentences [13, 14].

These great advancements allow for a two-step approach, where in the first step neural signals are transformed into a corresponding textual representations via a speech recognition system and in the second step this text is transformed into audible speech via text-to-speech synthesis.

However, using the textual representation as pivot has several disadvantages: (1) recognition / classification errors are propagated to the downstream processes, i.e. if the speech recognition system produces wrong results, the speech synthesis will be wrong as well, (2) since the recognition process needs to take place prior to resynthesis, the two-step approach is too time-consuming for real-time scenarios where immediate audio feedback is needed, and (3) speech carries information about prosody, emphasis, emotion etc. which are lost once transformed into text. For these reasons, it would be desirable for spoken communication to directly synthesize an audible speech waveform from the neural recordings [15].

In [16], Santoro et al. reconstructed the spectro-temporal modulations of real-life sounds from fMRI response patterns. Kubanek et al. [17] were able to track the speech envelope from ECoG. Neural signatures of speech prosody in receptive [18] and productive [19] speech cortices have been described. Studies were able to reconstruct perceived speech from ECoG signals [20] and to reconstruct spectral dynamics of speech from ECoG [21]. In a pilot study, we showed that it was possible to reconstruct an audio waveform from ECoG signals during speech production [22], but to the best of our knowledge, no study has reconstructed a high-quality audio waveform of produced speech from ECoG using deep neural networks.

Neural networks have shown great success in speech recognition [23] and speech synthesis [24]. However, so far they have been used only rarely for brain recordings. This is partially due to the very limited amount of speech data available for individual participants and the requirement to train speaker dependent models. In traditional speech processing, speaker dependent systems use tens of hours of data for training while high performing speaker independent systems accumulate thousands of hours.

More recently, initial studies have successfully applied deep learning methods to brain data [25, 26, 27] and BCI applications [28, 29, 30, 31, 32, 33]. Here, we show that densely-connected convolutional neural networks can be trained on limited training data to map ECoG dynamics directly to a speech spectrogram. This densely connected network architecture is specifically tailored to cope with the small amount of data. We then use a Wavenet vocoder [34] conditioned on logMel features to transform the reconstructed logMel spectrogram to an audio waveform. In this study, we restrict the vocoder to the conversion from an encoded speech representation to an audible wavefile. The Wavenet vocoder uses a much larger training corpus to map the logMel spectrograms directly onto an acoustic speech signal. The resulting audio is of high quality and reconstructed words are often intelligible. Thus, we present the first deep neural network reconstructing high-quality audio from neural signals during speech production.

## 2. Material and Methods

### 2.1. Experiment and Data Recording

We recorded ECoG from six native English speaking participants while they underwent awake craniotomies for brain tumor resection. All participants gave informed consent and the study was approved by the Institutional Review Board at Northwestern University. All subjects had normal speech and language function and normal hearing. ECoG was recorded with a medium-density, 64-channel, 8 × 8 electrode grid (Integra, 4 mm spacing) placed over the ventral motor cortex (M1v), premotor cortex (PMv) and inferior frontal gyrus pars opercularis (IFG). Grid locations were determined using anatomical landmarks and direct cortical stimulation to confirm coverage of speech articulatory sites. ECoG recordings were sampled at 2 kHz with Neuroport data acquisition system (Blackrock Microsystems, Inc.).

Participants were asked to read aloud single words shown to them on a screen while we recorded ECoG signals. These words were predominantly monosyllabic and consonant-vowel-consonant, and mostly compiled from the Modified Rhyme Test [35]. Participants read between 244 and 372 words resulting in recording length between 8.5 and 12.7 minutes. Note the extremely limited amount of subject dependent data compared to traditional speech synthesis. Acoustic speech was recorded using a unidirectional lapel microphone (Sennheiser) and sampled at 48 kHz. Figure 1 visualizes our experimental procedure. Stimulus presentation and simultaneous recording were facilitated using BCI2000 [36].

**Figure 1.**
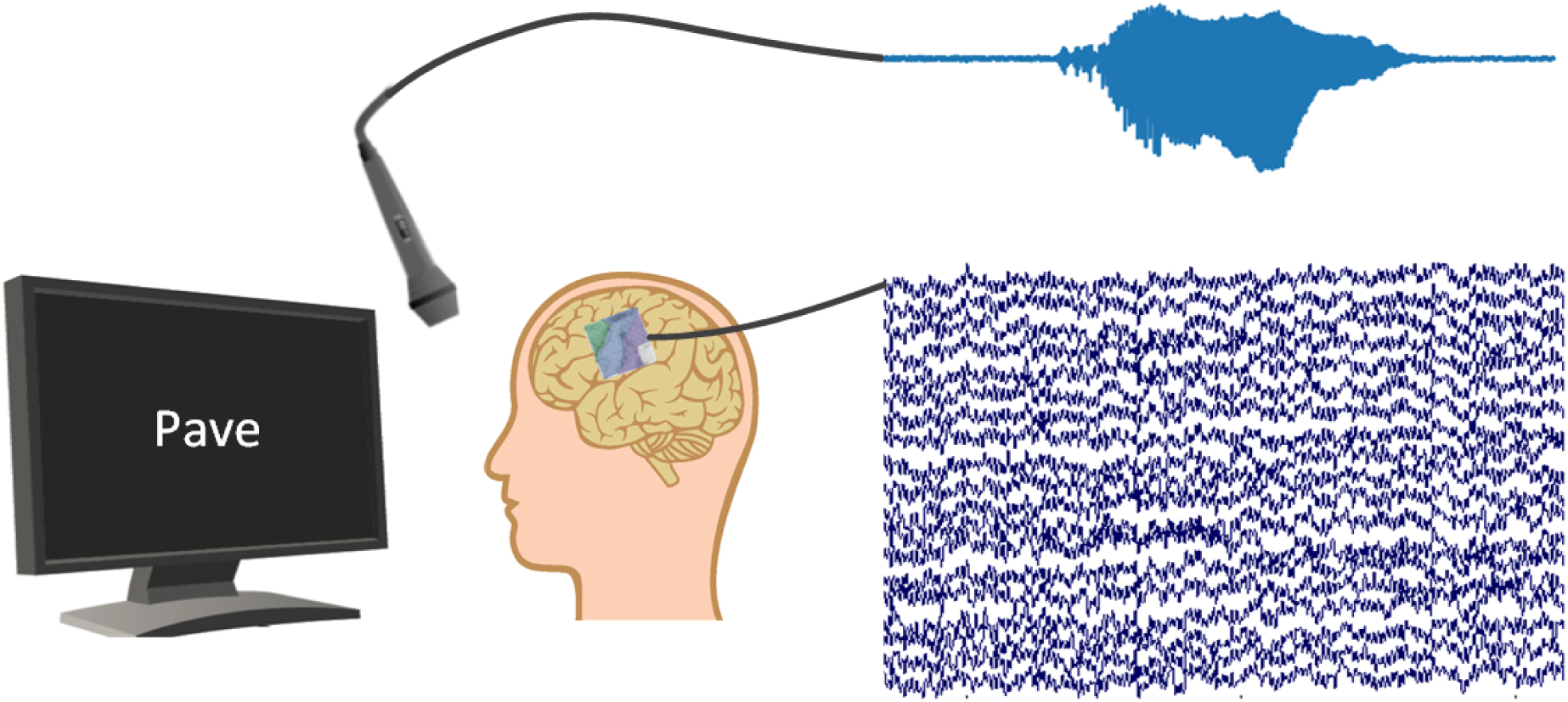
Illustration of the experiment. Participants are asked to repeat words shown on a screen. During speech production, ECoG data and acoustic stream are recorded simultaneously.

### 2.2. Data Processing

We first applied linear detrending to the ECoG signals and attenuated the first harmonic of the 60 Hz line noise using elliptic IIR notch filter. To extract meaningful information from the ECoG signals, we calculated logarithmic power in the broadband-gamma frequency range (70-170 Hz), which is known to largely reflect ensemble spiking [37] and contains the most amount of information about movement and speech processes [38]. Broadband-gamma power was extracted in 50 ms windows with an overlap of 10 ms. This time interval was chosen short enough to capture the fast processes of speech production while simultaneously being long enough to estimate broadband-gamma reliably. To estimate signal power, we calculated the mean of the squared signal in each window. We applied a logarithm transform to the extracted broadband-gamma features to make their distribution more Gaussian.

We normalized broadband-gamma activity of each electrode individually to zero mean and unit variance. To capture the long-range dependencies of the speech production process [39, 40], we included neighboring broadband-gamma activity up to 200 ms into the past and future in our study, resulting in 9 temporal contexts being stacked to form our final features. This results in a feature space of size 64 *electrodes ×* 9 *temporal offsets*. To represent the spatial topography of the electrode array, we arranged our features as a 8 × 8 × 9 matrix for decoding each time window.

For processing the acoustic data, we first downsampled the speech signal to 16 kHz. We transformed the waveform data into the spectral domain using 50 ms windows with an overlap of 10 ms to maintain alignment with the neural data. We discarded the phase information and only used the magnitude of the spectrograms [21, 41, 20]. To compress the magnitude spectrograms, we used 40 logarithmic mel-scaled spectral bins which should better represent the acoustic information in the speech data [42]. The logarithmic mel-scaled spectrograms (logMels) are extracted by taking the magnitude spectrogram and mapping it onto the mel-scale using filter banks. From now on we refer to the logMel representation as the spectrogram.

### 2.3. Decoding Approach

To transform the recorded neural signals into an audio waveform, we first trained densely connected convolutional neural networks (DenseNets) [43] to map the spatio-temporal broadband-gamma activity onto logarithmic mel-scaled spectrograms. This was done for each participant individually, as electrode grid placements and underlying brain topologies are vastly different between participants. The DenseNet regression model is described in Section 2.4.

We then used a Wavenet vocoder conditioned on the same spectral features of speech to recreate an audio waveform from the reconstructions of our densely connected convolutional neural network. For this Wavenet vocoder, a much larger data corpus could be used, as no user specific mapping had to be learned. Section 2.5 describes the Wavenet vocoder in more detail. Figure 2 highlights our decoding approach. The broadband-gamma activity over time (purple) is fed into the DenseNet regression model to reconstruct spectral features of speech (yellow). These are then transformed into an audio waveform using the Wavenet vocoder.

**Figure 2.**
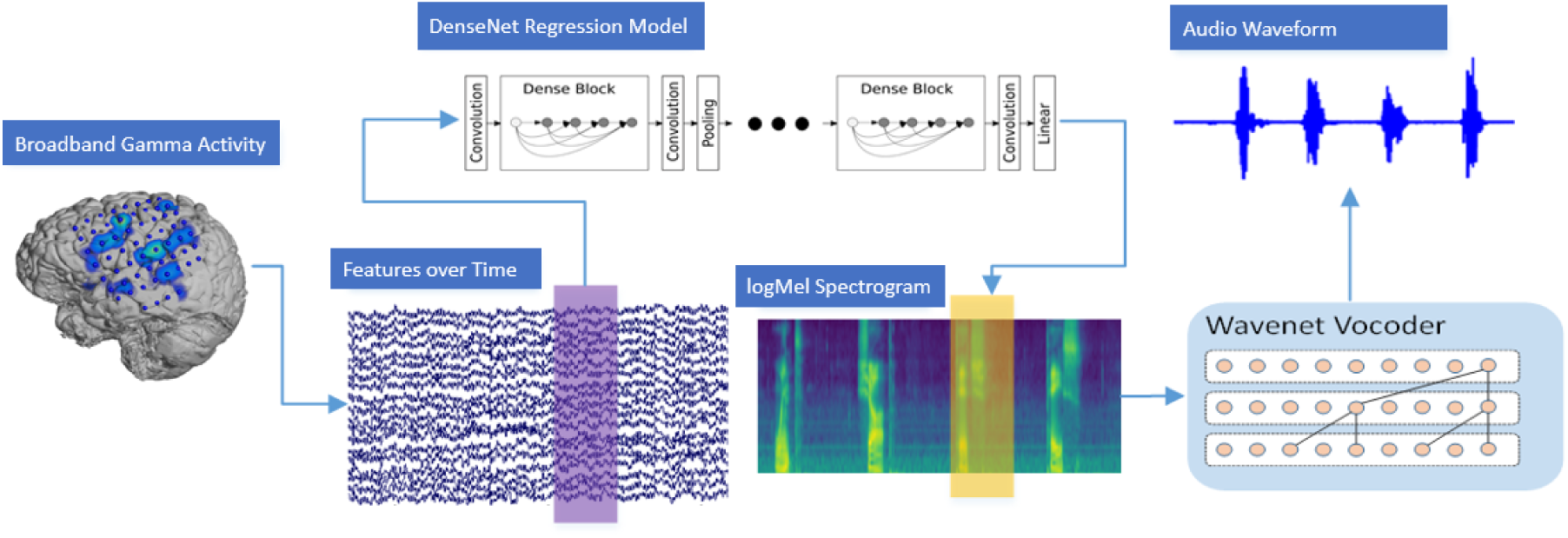
Overview of the decoding approach illustrating the transformation of neural data into an audible waveform. ECoG features for each time window are fed into DenseNet regression model to reconstruct the logarithmic mel-scaled spectrogram. Wavenet is then used to reconstruct an audio waveform from the spectrogram.

### 2.4. DenseNet Regression Model

The DenseNet architecture is a feed-forward multilayer network which uses additional paths between earlier and later layers in a dense structure. Each layer receives all feature maps of previous layers as an input and propagates its own feature maps to all subsequent layers. By passing the feature maps to subsequent layers, shortcut connections are established which improve the gradient and information flow in the training. This behavior is similar to the ResNet[44] architecture, where an identity function is used to circumvent the degradation problem. DenseNets were previously found to require fewer parameters to achieve a similar performance as ResNet [43]. ResNet’s solution relies on a residual mapping which combines the input and the output of a layer sequence by an addition operation. In contrast, DenseNet concatenates the feature maps of preceding layers. This way, each convolution adds its local information to the collective knowledge in the network and therewith forms an implicit global state. In a classification task, the final softmax layer can be seen as a classifier which takes the output of the previous layer and all preceding feature maps under consideration for its prediction.

A few adjustments needed to be done to adapt DenseNet for our regression task. On the one hand, it is intended for the convolution operations to apply on all three dimensions, namely the *x* and *y* position within the electrode grid and *time*, to find pattern in the spatio-temporal space. We therefore used 3D convolutions as well as 3D pooling layers throughout the network instead of their 2D counterparts used in traditional image processing. On the other hand, we changed the output layer to a fully connected layer with 40 neurons and a linear activation function to create a continuous output for the spectral coefficients.

Figure 3 shows an overview of the network structure and its integration inside the synthesis pipeline. The architecture consists of three dense blocks which group together a sequence of sublayers. Each sublayer is a composition of the following consecutive functions: Batch Normalization, rectified linear unit (ReLU) and a 3×3×3 convolution. The number of feature maps is set initially to 20 and increases according to a growth rate of *k* = 10. Two sublayers were used in each dense block yielding a total amount of 80 feature maps. Dense blocks are connected through transition layers, which are used as a downsampling operation for the feature maps of preceding layers, to fit their dimensions for the input of the next block. The output layer is a regressor, which estimates the spectral coefficients based on the input data and the feature maps in the collective knowledge. Overall, our resulting network comprises around 83,000 trainable parameters. For the training procedure, we used Adam[45] to minimize the mean squared error loss and used a fixed number of 80 epochs.

**Figure 3.**
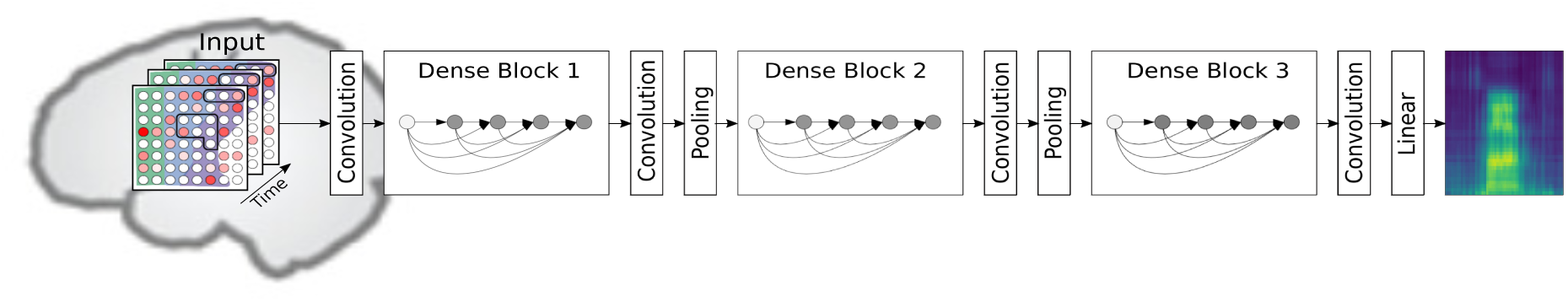
Overview of the DenseNet network structure. Input samples are preprocessed features of the neural signal with the shape 8 × 8 × 9. The first two dimensions are used for the spatial alignment of the electrodes, while the third dimension comprises the temporal dynamics. The network architecture consists of three Dense Blocks to map the neural features onto the speech spectrogram.

### 2.5. Wavenet Vocoder for the Reconstruction of Audible Waveforms

In the transformation process of acoustic data into logarithmic mel-scaled coefficients, we discard phase information. Nevertheless, this information is crucial for the inversion from the frequency domain back into a temporal signal. Recent studies show that high quality speech waveforms can be synthesized by using Wavenet [46] conditioned on acoustic features estimated from a mel-cepstrum vocoder [47]. During network training, the model learns the link between speech signal and its acoustic features automatically without making any assumptions about prior knowledge of speech. The synthesized acoustic waveform generated by Wavenet recovers the phase information previously lost.

In this paper, we conditioned Wavenet in the same way on logarithmic mel-scaled coefficients as described in Tacotron-2 [34] for Text-to-Speech synthesis. The filterbank features tend to outperform cepstral coefficients for local conditioning [48]. We used a separate dataset for training our Wavenet model. The LJ-Speech corpus [49] contains utterances from a single speaker reading passages from seven non-fictional books. The length of an utterance varies between one and ten seconds and sums up to a total amount of around 24h of speech data. Empirical tests based on the spectral coefficients of the reference data showed that this corpus is suitable to train Wavenet and reconstruct high quality acoustic waveforms containing intelligible speech and capturing the speaker characteristics of the participants.

The internal architecture is depicted in Figure 4. The model expects two feature matrices with possibly differing dimensions as inputs: the acoustic speech waveform **x** and the spectral features **c** for the local conditioning. Due to the fact that the dimensions of both input data might not match, a transformation is needed as an adjustment. An initial 1 × 1 convolution is used to increase the number of channels, known as residual channels. The spectral features get upsampled by four consecutive transpose convolutions to match the dimensions with the convolved acoustic speech signal.

**Figure 4.**
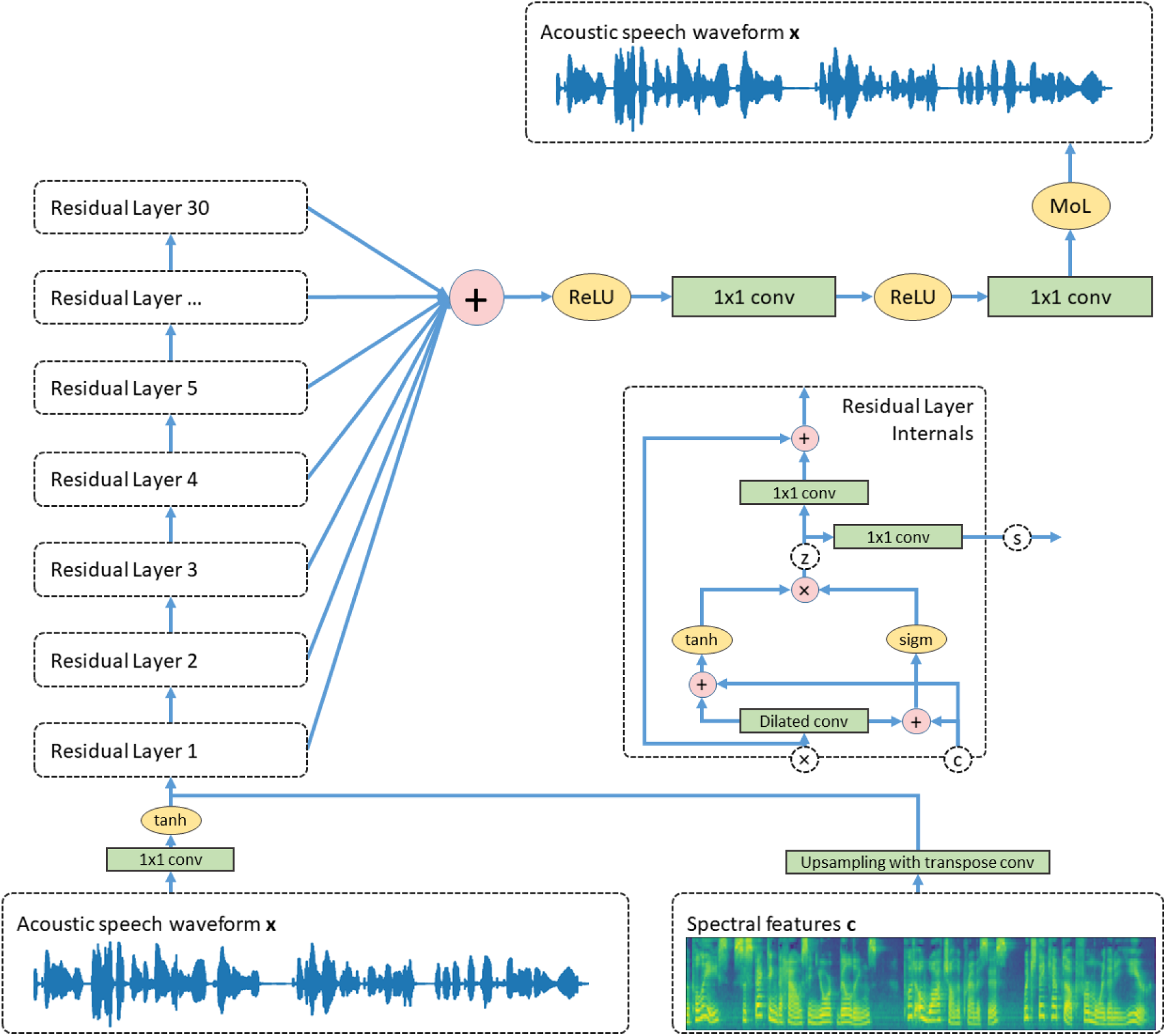
Overview of the Wavenet vocoder architecture. The network comprises a stack of 30 residual blocks to find a mapping between the acoustic speech signal **x** to itself considering the extracted features **c**. Each block has a separate output which are summed in the calculation of the actual prediction. We use a 10-component mixture of logistic distributions (MoL) for the prediction of audio samples.

After adjusting both input sequences, a stack of residual blocks is used whose interior is illustrated inside Figure 4. Each block contains a gated activation function which calculates a hidden state *z* given the following equations:

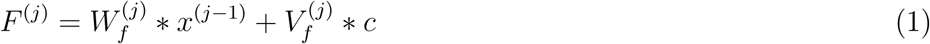

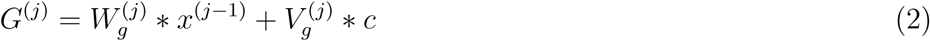

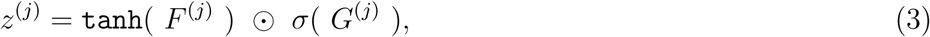

where * implies a dilated causal convolution, ⊙ denotes the hadamard product and *σ*(·) corresponds to the sigmoid function. The superscript *j* indicates the current residual block. *W_f_*, *W_g_*, *V_f_* and *V_g_* are trainable weights of the convolution. In the gated activation function, the equations *F* and *G* indicate the filter and gate, respectively. A residual block uses two outputs to speed up the training procedure of the network. Both outputs are based on the intermediate result of the hidden state *z*. The residual blocks are connected throughout their stack by using residual connections [44] which enable training of very deep neural networks. For the prediction of the next audio sample, the network uses skip connections. Both outputs are computed in the following way:

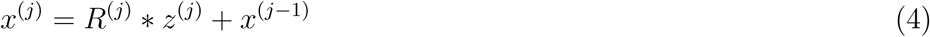

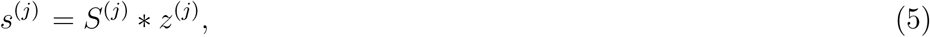

where * denotes a 1 × 1 convolution to adjust the dimensionality of the channels. *R* and *S* represent trainable weights of their convolution operation.

In the stack of residual blocks, the dilation rate of the dilated causal convolution increases exponentially and gets reset after a fixed amounts of layers to start a new cycle. Parameter choices have been made in accordance to the Tacotron-2 System [34] which results in a stack of 30 residual blocks and three dilation cycles.

Wavenet is an autoregressive model which predicts the next audio sample based on all previously seen samples inside its receptive field and its conditioning features:

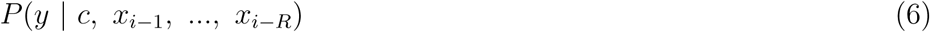

For the prediction of the next audio sample, the model considers all skip connections from the residual blocks by summation and processes the results through a sequence of rectified linear units and 1 × 1 convolutions as shown in Figure 4.

Recent improvements suggest to model the generated speech with a discretized mixture of logistic distributions [50, 51]. We follow the proposed approach from Tacotron 2 and use a linear projection layer to predict the parameters for each mixture component [34]. The Wavenet vocoder uses a 10-component mixture of logistic distributions to model the reconstructed speech signal.

In network training, we use Adam as our optimization method with an initial leaning rate of 0.001. We trained our Wavenet vocoder for a fixed amount of 600,000 update steps. All hyperparameter choices are summarized in Table 1. We based our Wavenet vocoder on an open source implementation available on GitHub [52].

**Table 1.**
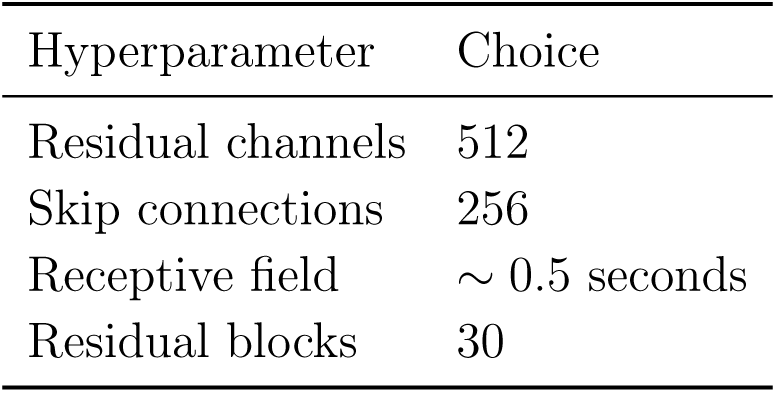
Overview of the hyperparameter choices made for our Wavenet vocoder.

| Hyperparameter | Choice |
| --- | --- |
| Residual channels | 512 |
| Skip connections | 256 |
| Receptive field | $\sim 0.5$ seconds |
| Residual blocks | 30 |

## 3. Results

For each participant, we partitioned the recorded data into disjoint sets for training and testing in a 5-fold cross validation. This approach ensured that the complete spectrogram could be reconstructed without operating on data that has been seen before. Twenty percent of the training data are reserved as a separate validation set to analyze the optimization process of the network training. The reconstruction from DenseNet is frame based and yields a spectrogram of the same length as the original one.

We used the Pearson product-moment correlation for the evaluation of our synthesized speech. The correlations are computed for each frequency bin individually between the reconstructed spectrogram and its original counterpart. For comparison, we computed a chance level by splitting the data at a random point in time and swapping both partitions. This ensures that the structure of the neural signals is preserved but the alignment to the spectral representation of speech is shifted. To reconstruct a spectrogram based on broken alignment, we performed additional network training in a 5-fold cross validation with the randomized dataset. For each participant, we repeated the estimation of a random chance level 20 times by using unique splits along the temporal dimension to approximate chance level.

Figure 5 (a) summarizes our results for all participants showing average correlations and their corresponding standard deviation. We achieve correlations significantly better than chance level (p<0.001) for all six participants with scores (mean ± standard deviation) of *r*_1_ = 0.19 ± 0.12, *r*_2_ = 0.29 ± 0.06, *r*_3_ = 0.56 ± 0.18, *r*_4_ = 0.34 ± 0.12, *r*_5_ = 0.69 ± 0.10 and *r*_6_ = 0.41 ± 0.07, respectively. Participant 5 clearly outperforms the other participants. We hypothesize that this might be due to better electrode placements.

**Figure 5.**
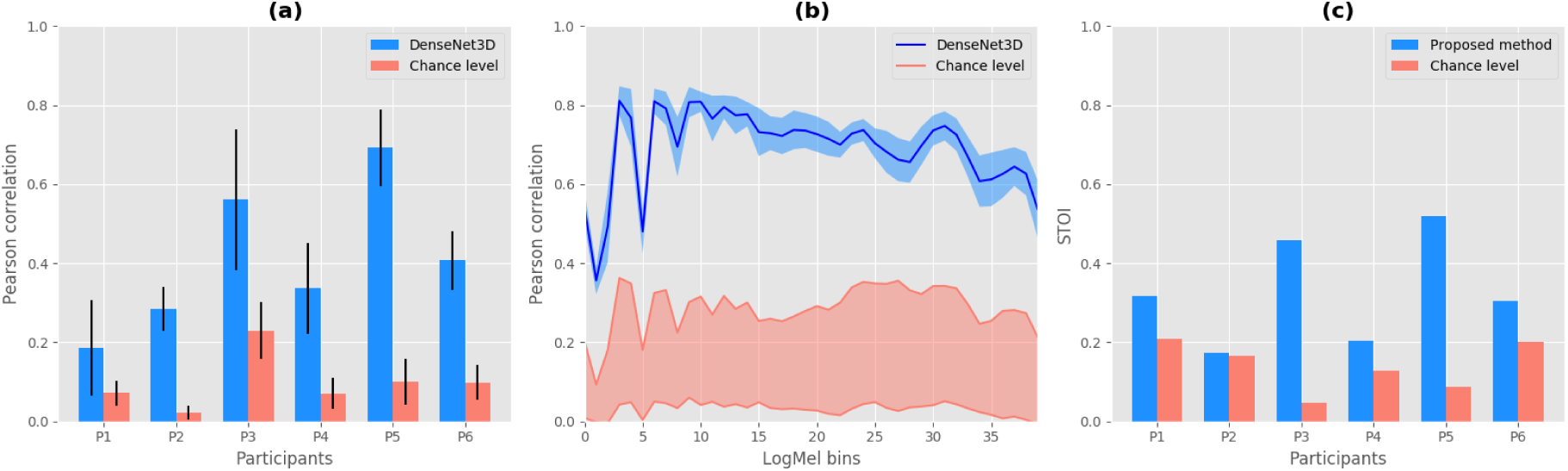
Reconstruction performance of DenseNet compared to random chance. (a) Pearson correlation coefficients between original and reconstructed spectrograms for each participant. Bars indicate the mean over all logarithmic mel-scaled coefficients while whiskers denote the standard deviation. (b) Detailed performance across all spectral bins for participant 5. (c) STOI scores as an objective intelligibility measure in comparison to the chance level.

Across all participants, correlations of reconstructed spectrograms are considerably above chance level.

In Figure 5 (b) we investigate the distribution of Pearson correlation coefficients for Participant 5 in more detail. For each spectral bin, the chance level is given by the minimum and maximum correlation coefficient under consideration of all 20 spectrograms from the baseline approximation. Our decoding approach achieves consistently high correlations above chance level across all spectral bins.

In order to evaluate the intelligibility of our reconstructed speech waveform, we employed the Short-Time Objective Intelligibilty measure (STOI) [53]. We used the original audio from the recording of the experiment as our reference. Figure 5 (c) reports the STOI scores for each participant. For the chance level, we took the spectrogram with the highest mean correlation under consideration to estimate a waveform using the Wavenet vocoder.

A reconstruction example of an excerpt from the experiment session of participant 5 is shown in Figure 6 (a) for visual inspection. The top row corresponds to the spectrogram of the reference data while the bottom row contains the time aligned spectral coefficients estimated by the DenseNet model. It is evident that the model has learned a distinguishable representation between silence and acoustic speech and captures many of the intricate dynamics of human speech. Furthermore, early characteristics of resonance frequencies are present in the spectral coefficients of the predicted word articulation.

**Figure 6.**
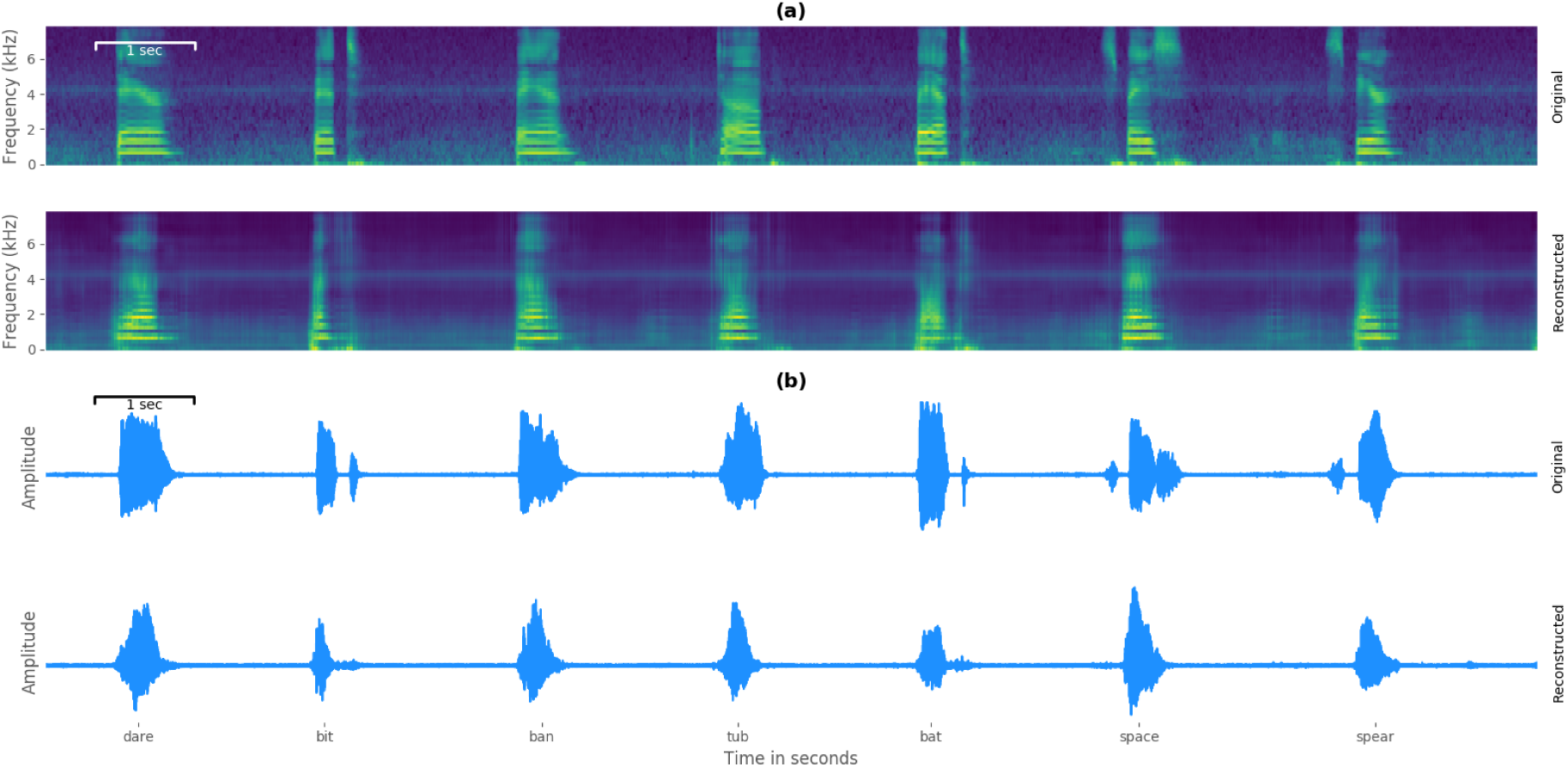
Reconstruction example for visual inspection. a) compares a time-aligned excerpt in the spectral domain of participant 5 and emphasizes the quality of the reconstructed acoustic speech characteristics. b) shows the generated waveform representation of the same excerpt as in the spectrogram comparison. Spoken words are given below.

Figure 6 (b) shows the resynthesized acoustic waveforms of the same excerpt from the reconstruction example by using the described Wavenet vocoder. Additionally, some listening examples of original and reconstructed audio can be found in the supplementary material. To compensate for artifacts of this conversion, we applied the transformation on the original and reconstructed spectrograms to isolate the synthesis quality of the trained network.

## 4. Discussion

We have shown that speech audio can be decoded from ECoG signals recorded from brain areas associated with speech production. We achieve this by combining two deep neural network topologies specifically designed for very different goals. In the first step, we employ a densely connected convolutional neural network which is well-suited to be trained on the extremely limited datasets. This network transforms the measured brain activity to spectral features of speech. Reasonable high correlations of up to *r* = 0.69 across all frequency bands were achieved by this network. Subsequently, a Wavenet vocoder is utilized to map these spectral features of speech back onto an audio waveform. As this model does not have to be trained for each participant individually, it is trained on a much larger speech dataset and the topology is tailored to maximize speech output quality. Selected examples of reconstructed audio can be found in the supplementary material where the original articulation of participant 5 and the corresponding reconstruction is presented in pairs.

To the best of our knowledge, this is the first time that high quality audio of speech has been reconstructed from neural recordings of speech production using deep neural networks. This is especially impressive considering the very small amount of training data available. In general, traditional speaker dependent speech processing models are trained with 10th of hours of data. This is an important step towards neural speech prostheses for speech impaired users. While our study uses overtly produced speech in non-paralyzed subjects, recent studis show analogous results for paralyzed patients [54, 55, 56, 57], as well as for speech decoding [58], providing reason to believe that similiar results might also be achievable for mute users.

## Supporting information

## Acknowledgment

MA, CH, DK and TS acknowledge funding by BMBF (01GQ1602) and NSF (1608140) as part of the NSF/NIH/BMBF Collaborative Research in Computational Neuroscience Program. MS acknowledges funding by the Doris Duke Charitable Foundation (Clinical Scientist Development Award, grant 2011039), a Northwestern Memorial Foundation Dixon Translational Research Award (including partial funding from NIH National Center for Advancing Translational Sciences, UL1TR000150 and UL1TR001422), NIH grants F32DC015708 and R01NS094748.

